# Distinct Motor Map Characteristics for Biceps and Triceps Muscles in Persons with Chronic Tetraplegia: Implications for Arm Function

**DOI:** 10.1101/2024.03.27.585546

**Authors:** Jia Liu, Kyle O’laughlin, Gail F. Forrest, Tarun Arora, Gregory Nemunaitis, David Cunningham, Steven Kirshblum, Svetlana Pundik, Kelsey Baker, Anne Bryden, Kevin Kilgore, Francois Bethoux, Xiaofeng Wang, M. Kristi Henzel, Nabila Brihmat, Mehmed Bugrahan Bayram, Ela B. Plow

## Abstract

Following spinal cord injury (SCI), intact neural resources undergo widespread reorganization within the brain. Animal models reveal motor cortical representations devoted to spared muscles above injury expand at the expense of territories occupied by weaker muscles. In this study, we investigated whether motor representations are similarly reorganized between a relatively spared biceps muscle and a weakened triceps muscle in persons with chronic tetraplegia following traumatic cervical SCI in association with upper limb motor function. Twenty-four adults with cervical SCI and 15 able-bodied participants underwent motor mapping using transcranial magnetic stimulation. We determined following map characteristics: area, amplitude (maximal motor evoked potential and volume), and center of gravity. Maximal voluntary contraction (MVC) and motor function (Capabilities of the Upper Extremity Test or CUE-T) were also assessed. Findings reveal that participants with SCI had hyper-excitable biceps maps than triceps, and hyper-excitable biceps maps also compared to biceps maps in able-bodied participants. Higher amplitude of biceps and triceps maps was associated with better motor function (higher CUE-T) and more distal injury (i.e., more spared segments) in persons with SCI. Amplitudes of biceps but not the triceps maps were associated with higher muscle MVCs. In conclusion, over-excitable biceps than triceps map in SCI may represent deafferentation plasticity. For the first time, we demonstrate how map reorganization of spared and weaker muscles in persons with chronic cervical SCI is associated with upper limb motor status. Use-dependent mechanisms may shift neural balance in favor of spared muscles, supporting potential use as response biomarkers in rehabilitation studies.

**New & Noteworthy:** Our study reports evidence in humans with cervical SCI that motor representation for the relatively spared muscle becomes hyper-excitable compared to that for the weaker muscle to the extent that hyper-excitability is even higher compared to biceps maps in uninjured individuals. Use-dependent mechanisms likely favor such heightened excitability of spared maps. For the first time, we demonstrate clinical relevance of map excitability in humans with SCI, supporting potential use as a biomarker of recovery.

## Introduction

Spinal cord injury is a leading cause of paralysis, affecting approximately 302,000 individuals in the United States (U.S.) alone [1]. Cervical spinal cord injury (SCI) is the most common (60%) and severe type of SCI [1]. It leads to characteristic paralysis of trunk and all four limbs, a clinical condition known as tetraplegia [1]. Weakness of the upper limbs limits independence to engage in activities of daily living (ADLs), self-care, and vocation and leisure [2]. Persons with tetraplegia therefore place a high priority on the return of upper limb motor function [2]. However, rehabilitation of upper limb function in tetraplegia remains limited.

Post-injury neural reorganization may be a factor important for functional recovery [3]. Following SCI, intact neural structures within the brain undergo widespread reorganization. Animal models reveal that motor cortical representations devoted to spared muscles above injury expand and become to take over cortical territories previously occupied by weaker muscles innervated below the injury [3–5]. Mechanisms of such deafferentation-plasticity likely involve unmasking of latent intracortical connections [6] and use-dependent plasticity [3] that evokes a “competition” for surviving resources amongst muscles with different extents of sparing during post-injury recovery.

Preliminary studies have indicated that similar post-injury reorganization may occur in humans with SCI investigated using non-invasive transcranial magnetic stimulation (TMS) [7, 8]. In a case series (n=2) [8] and a separate exploratory cross-sectional study (n=10) [7], investigators demonstrated that TMS-based motor cortical maps for the biceps muscle (which is generally stronger in typical cervical SCI) were larger and more excitable than maps for the weaker triceps muscle in persons with chronic cervical SCI. However, motor neurophysiology in humans with SCI can be influenced by heterogeneity of injury levels, motor impairment severity state, extent and type of rehabilitation received, etc. Therefore, a larger formative evaluation is necessary to validate such neural reorganization in humans with SCI and, importantly, to examine the clinical implications for upper limb motor function.

The purpose of this study was to investigate TMS motor cortical map characteristics for biceps and triceps muscles in individuals with tetraplegia following chronic traumatic cervical SCI. Further, we sought to examine how motor cortical map characteristics for these muscles were associated with upper limb function. We hypothesized that motor maps devoted to the stronger biceps muscle would be larger and more excitable than motor maps devoted to the weaker triceps muscle. We also anticipated that these map features would be associated with upper limb motor function, setting the stage for map characteristics as targets for designing effective rehabilitative treatments in SCI.

## Methods

### Participants

Individuals aged 18-75 years with chronic (>1 year) traumatic cervical SCI (C1-C8) were included. Participants could have complete or incomplete injuries (defined as American Spinal Injury Association Impairment Scale (AIS) A-D) as long as the following minimal motor criteria were met in the more affected arm: a stronger biceps muscle, rated as 3-5 (inclusive) on the Medical Research Council (MRC) scale, and a weaker triceps muscle, rated ≥1 MRC grade lower than biceps MRC (i.e., 1-3 MRC, inclusive). (Note that according to the MRC scale: 0=no movement or contraction; 1=palpable or visible twitch/contraction; 2=active movement, full range of motion, gravity eliminated; 3=active movement, full range of motion, against gravity; 4=active movement, full range of motion, against gravity and some resistance; and 5=active movement, full range of motion, against gravity and maximal resistance). Persons with AIS A, i.e., “complete” injury were included because specified motor criteria would allow for zones of partial preservation in the innervation of biceps and triceps muscles. Bilateral arms were assessed, and the more affected arm was identified as the one that possessed a lower triceps MRC grade. Licensed physical/occupational therapists evaluated muscle grades to determine eligibility, and a board-certified physician confirmed the SCI diagnosis and AIS grade.

We also enrolled able-bodied individuals without neurological or musculoskeletal conditions. People with contraindications to TMS (e.g., pacemaker, metal in the skull, history of seizures, pregnancy) were excluded [9]. All participants (**Table 1**) were part of a multi-site clinical trial investigating effects of transcranial direct current stimulation paired with rehabilitation in cervical SCI (Cleveland Clinic, Cleveland Veteran Affairs, and Cleveland MetroHealth in Ohio, and Kessler Foundation in New Jersey; ClinicalTrials.gov identifier: NCT03892746) [10]. Baseline cross-sectional data are presented here for this study. All participants gave a written informed consent approved by the site-specific Institutional Review Board and the Human Research Protections Office of the U.S. Department of Defense.

**Table 1.** Participant demographics and SCI characteristics.

| Group | ID | Sex | Age (year) | Months Since SCI | AIS | NLI | ML | SL | More Affected Arm |  |  |
| --- | --- | --- | --- | --- | --- | --- | --- | --- | --- | --- | --- |
|  |  |  |  |  |  |  |  |  | Triceps MRC | Biceps MRC | UEMS (/25) |
| SCI | 01 | M | 32 | 20 | D | C6 | C6 | T14 | 3 | 5 | 15 |
|  | 02 | M | 61 | 18 | C | C5 | C6 | C5 | 3 | 5 | 11 |
|  | 03 | M | 23 | 36 | B | C4 | C5 | C4 | 2 | 4 | 8 |
|  | 04 | M | 32 | 108 | B | C5 | C6 | C5 | 2 | 5 | 12 |
|  | 05 | M | 46 | 161 | D | C4 | C5 | C4 | 3 | 4 | 18 |
|  | 06 | M | 59 | 137 | D | C2 | C6 | C2 | 2 | 4 | 17 |
|  | 07 | M | 62 | 108 | D | C4 | C5 | C4 | 3 | 4 | 16 |
|  | 08 | F | 31 | 90 | B | C5 | C5 | C5 | 1 | 4 | 6 |
|  | 09 | M | 55 | 36 | D | C7 | C7 | T2 | 3 | 5 | 22 |
|  | 10 | M | 53 | 402 | D | C5 | C5 | C5 | 3 | 4 | 16 |
|  | 11 | M | 47 | 48 | A | C6 | C6 | C6 | 2 | 4 | 10 |
|  | 12 | M | 63 | 242 | D | C8 | C8 | C8 | 3 | 5 | 18 |
|  | 13 | F | 35 | 33 | C | C3 | C8 | C4 | 3 | 4 | 17 |
|  | 14 | M | 29 | 120 | B | C6 | C6 | C8 | 3 | 4 | 12 |
|  | 15 | M | 46 | 45 | D | C2 | C2 | C2 | 2 | 3 | 10 |
|  | 16 | M | 63 | 29 | D | C4 | C5 | C4 | 2 | 3 | 11 |
|  | 17 | M | 72 | 24 | C | C2 | C5 | C4 | 2 | 4 | 9 |
|  | 18 | M | 49 | 14 | A | C6 | C6 | C6 | 3 | 5 | 13 |
|  | 19 | F | 45 | 20 | D | C4 | C7 | C4 | 3 | 4 | 15 |
|  | 20 | F | 29 | 89 | A | C4 | C5 | C4 | 2 | 4 | 6 |
|  | 21 | M | 74 | 15 | C | C1 | C1 | C1 | 1 | 3 | 9 |
|  | 22 | M | 69 | 107 | D | C3 | C3 | C3 | 2 | 4 | 14 |
|  | 23 | M | 64 | 180 | A | C4 | C6 | C6 | 3 | 5 | 13 |
|  | 24 | M | 42 | 276 | B | C5 | C6 | C6 | 1 | 4 | 9 |
|  | n=24 | 4 (F)<br>20(M) | 49.2<br>(15.2)<br>* | 98.5<br>(97.3) | 4(A)<br>5(B)<br>4(C)<br>11(D) | 13(C1-4)<br>11(C5-8) | - | - | 3<br>(2, 3)<br>a | 4<br>(4, 5)<br>a | 12.8<br>(4.1) |
| Control | n=15 | 10(F)<br>5(M) | 35.8<br>(14.3)<br>* | - | - | - | - | - | - | - | - |

F=female; M=male; NLI=neurological level of injury; ML=motor level; SL=sensory level; AIS=American Spinal Injury Association impairment scale; MRC=Medical Research Council; UEMS=upper extremity muscle score. Data were mean (SD). \*: significant difference in age between SCI and control groups, $p<0.05$ . <sup>a</sup>: significant difference in MRC grades between biceps and triceps muscles, $p<0.001$ .

### Neurophysiological Assessments

Participants sat in a height-adjusted chair or wheelchair with shoulder in neutral or slight flexion/abduction and elbow in 90° flexion. Bi-polar Ag/AgCl electromyography (EMG) electrodes were placed in a belly-belly arrangement on biceps and triceps muscles of the more affected arm, with a ground electrode on the clavicle. Among able-bodied participants, we tested the dominant and non-dominant arm across alternating enrollments to eliminate the confounding effect of side of dominance (assessed using the Edinburgh Handedness Inventory [11]).

EMG data were recorded using PowerLab (AD Instruments, Colorado Springs, CO, USA) at a sampling rate of 4 kHz from 3 study centers (i.e., Cleveland Clinic, MetroHealth Center for Rehabilitation Research, and Kessler Foundation) and using Brain Vision (Brain Vision, Morrisville, NC, USA) at a sampling rate of 5 kHz from another center (i.e., Cleveland Veteran Affairs Medical Center, for 4 participants with SCI and 3 without SCI). A uniform high-pass filter at 10Hz was applied during data collection to eliminate movement-related artifacts. Neurophysiologic testing was standardized across research sites using video tutorials, images, and a written Standard Operating Procedure manual.

We first collected maximal voluntary contraction (MVC) activity generated by biceps and triceps muscles. Participants were instructed to exert maximal isometric forces against external resistance, which meant attempting to pull up supinated forearm towards ipsilateral shoulder for biceps, and to push down forearm in neutral rotation against rigid surface for triceps. Three, 5-second MVC trials were collected for each muscle, with ≥30 seconds of inter-trial interval to mitigate fatigue. We applied root-mean-square (RMS) calculation to each MVC trial using a 2-second smoothing window. The mean RMS value of middle 3 seconds of each MVC trial was calculated and then averaged across 3 trials of the same muscle. The averaged RMS value was defined as the MVC value for each muscle.

Next, TMS was applied using a Magstim 200^2^ device (Magstim, Dyfed, UK) which delivers monophasic pulses through a figure-of-eight coil (70mm diameter). The coil was held tangential to the scalp in the posterior-anterior direction with a 45° angle orientation to midline. The coil position was guided using a neuro-navigation system (Brainsight, Rogue Research, Montreal, QC, Canada). A template brain image was used to register cranial surface landmarks for each participant. Registration errors of no more than 6mm were permitted with visual verification on Brainsight software.

Prior to conducting TMS motor mapping, we determined motor cortical hotspots for the biceps and triceps muscles. TMS pulses were delivered to the motor cortex contralateral to the more affected arm, while participants maintained slight isometric contraction of the target muscle (20±5% MVC). The motor hotspot was defined as the site that elicited criterion-sized motor evoked potentials (MEPs) at the lowest TMS intensity in ≥ 6/10 trials [12, 13]. The criterion was defined as MEPs with peak-to-peak amplitude ≥ 0.1mV above pre-stimulus EMG activity (measured 150ms to 5ms before the TMS stimulus). The lowest TMS intensity used to elicit criterion-sized MEPs at hotspot was defined as the active motor threshold (AMT).

Subsequently, TMS mapping was performed across a 7-by-7 grid (1 cm^2^ resolution) placed over the motor cortex and centered at the biceps hotspot. Five TMS pulses were delivered at each of 49 scalp sites (at 100% maximum stimulator output or MSO), while both biceps and triceps muscles were at rest. This approach of using 100% MSO has also been used by Brouwer and Hopkins-Rosseel [7] and Levy Jr et al [8] and testing at rest allows examination of motor maps of two muscles at the same time without the confounding effect of differing intensities while minimizing fatigue. We monitored pre-stimulus (100ms to 10ms before TMS artifact) EMG activity during TMS mapping (cohort median [25^th^ and 75^th^ percentiles] pre-stimulus activity for biceps: 3.0 [2.1, 4.4] μV; for triceps 4.8 [2.7, 8.9] μV).

We used a customized MATLAB script to analyze data. Across 49 scalp locations, active motor sites were defined as those that elicited peak-to-peak MEP amplitudes ≥ 0.05mV in at least 3/5 trials. The mean MEP amplitude was calculated for each active site. Following TMS mapping features were calculated across active motor sites separately for biceps and triceps maps: area (i.e., total number of active sites, cm^2^), max-MEP amplitude (i.e., the largest mean MEP found across active sites, mV), volume (i.e., summed mean MEPs across active sites, mV*cm^2^), and center of gravity in the mediolateral direction (COG (x), equation 1) and in the anterior-posterior direction (COG(y), equation 2) [14]. In participants whose left hemisphere was targeted during TMS, the sign of COG (x) was reversed before statistical analyses. If the TMS maps could not be successfully elicited (i.e., zero active site), zeros were used for map area, max-MEP, and volume, and missing values were used for COG (x) and COG (y).

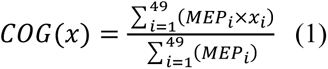

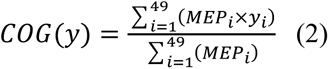

### Functional Assessments

Motor functional ability of the upper limbs was defined in participants with SCI using the Capabilities of Upper Extremity Test (CUE-T) which has excellent test-retest reliability (intraclass correlation coefficient = 0.97-0.98) and good-to-excellent concurrent validity (0.55-0.83) in SCI [15, 16]. CUE-T consists of 15 unilateral and two bilateral tasks which assess gross and fine motor abilities of the proximal and distal upper limb segments. Each task is scored 0, 1, 2, 3, or 4 by a trained therapist based on repetitions completed, weight manipulated/lifted, force generated, or time needed to complete an item. For the interest of this study, scores were summed from the following 7 items that require activation of triceps and biceps muscles of the more affected arm: *Reach Forward*, *Reach Up*, *Reach Down*, *Pull Weight*, *Push Weight*, *Lift Up* (bilateral), and *Push Down* (bilateral), where the total subscore can range from 0 to 28.

### Statistical Analysis

Two-way ANOVA was used to determine main and interaction effects of GROUP (SCI vs control) and MUSCLE (biceps vs triceps) on mapping variables of interest (i.e., area, max-MEP amplitude, volume, COG (x), and COG (y)). If there was a significant interaction effect, post-hoc analyses were conducted with Bonferroni correction for multiple comparisons. Depending on whether data were normally distributed (per Shapiro-Wilk tests), t-tests or Wilcoxon-Signed Ranks Sum tests were used.

To examine the relationship between TMS map characteristics and upper arm CUE-T scores, Pearson’s (r) or Spearman (ρ) correlation was performed, subject to results of normality tests. Given that tasks assessed using CUE-T (e.g., *Reach Forward*, *Reach Up*) mostly require sub-maximal efforts to complete, we also examined the correlations between TMS map characteristics and MVCs which corresponds to maximal effort. Exploratory correlation analyses were also performed between muscle MVCs and upper arm CUE-T scores and between TMS map characteristics and clinical injury severity (given by AIS grade), NLI, post-injury chronicity (in months), and severity of muscle spasticity (measured by the Modified Ashworth Scale or MAS). All statistical analyses were performed using R (version: 4.2.2) with a significance level of alpha = 0.05.

## Results

Twenty-four persons with SCI and 15 able-bodied individuals were studied (**Table 1**). Representative TMS maps and MVCs of biceps and triceps muscles are presented in **Fig. 1** comparing between an able-bodied participant and an individual with SCI (ID1, **Table 1**). Our results of muscle MVCs and AMTs confirmed differential sparing of biceps versus triceps muscles in participants with SCI. Compared to uninjured persons, participants with SCI had a marked reduction of triceps muscle MVCs (0.2±0.2 mV vs 0.4±0.2 mV, p<0.001) but not of biceps muscle MVCs (0.6±0.5 mV in SCI vs 0.7±0.4 mV in control, *p*=NS). Participants with SCI also had significantly higher AMTs for triceps (58.2±13.6 %MSO vs 45.6±7.4 %MSO, *p*<0.001) but not biceps (42.5±15.9 %MSO in SCI vs 41.2±6.8 %MSO in control, *p*=NS) muscles than able-bodied individuals. We successfully elicited biceps TMS maps in 22/24 (92%) participants with SCI and 14/15 (93%) able-bodied controls. Triceps maps were elicited in 21/24 (88%) of participants with SCI and 12/15 (80%) of able-bodied controls. CUE-T was tested in all except 1 participant with SCI who did not return after TMS assessments (ID23, **Table 1**).

**Figure 1.**
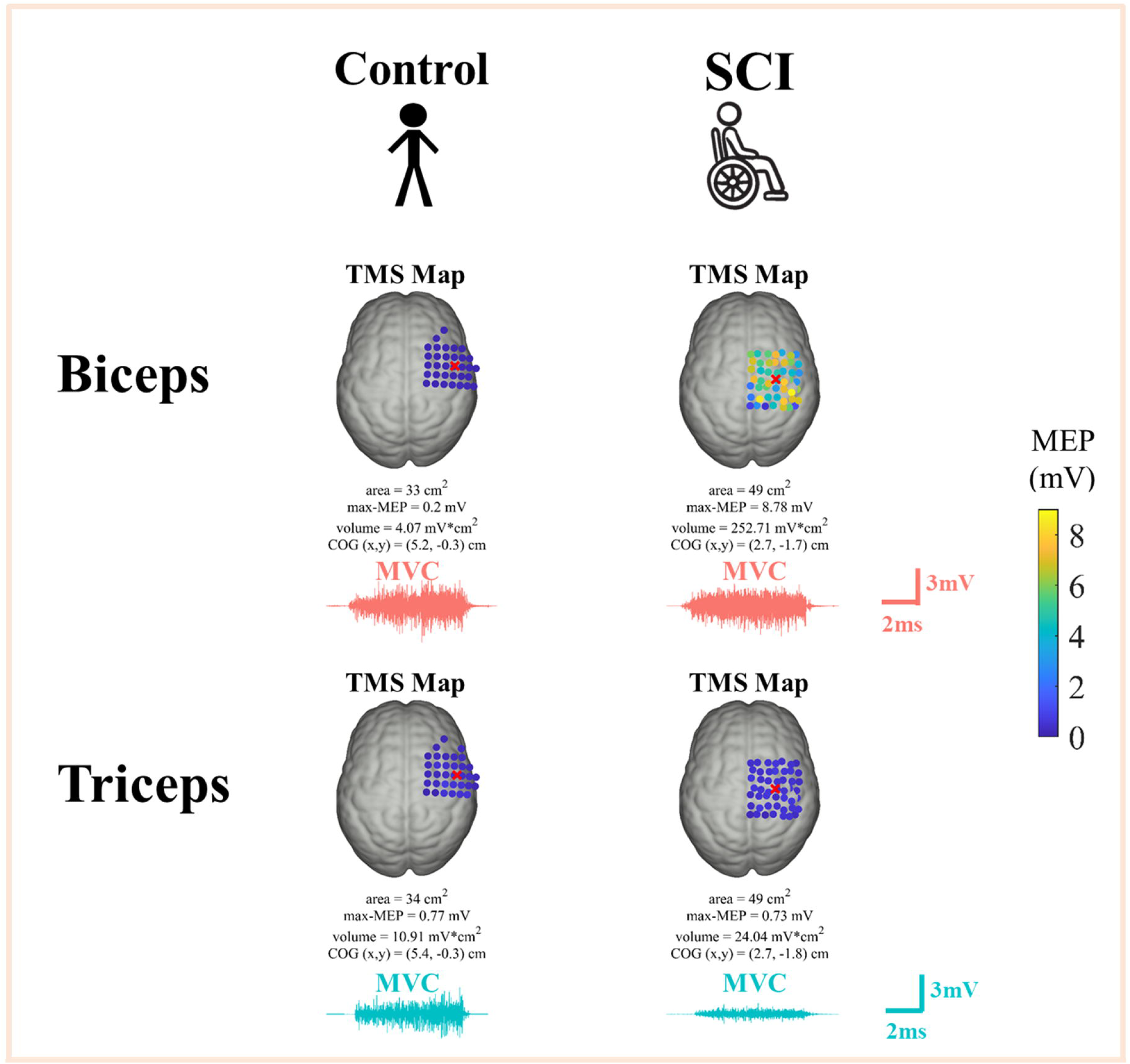
Representative TMS maps and muscle MVCs. Compared to the representative control participant (left), the individual with SCI (right) showed pronounced over-excitability in the biceps TMS map, although MVC values were comparable between two participants. With respect to the triceps muscle, TMS map excitability was similar between two participants, whereas there was a marked reduction of triceps MVC in the participant with SCI than the uninjured control. Red X marker in each TMS map indicates the location of COG.

### Map Characteristics

TMS map maximal excitability defined using max-MEP amplitudes showed significant main effects of GROUP (F_(1,37)_= 5.4, *p=*0.026) and MUSCLE (F_(1,37)_= 8.8, *p=*0.005) as well as an interaction effect (F_(1,37)_= 6.5, *p=*0.015, **Fig. 2A**). Posthoc analysis revealed that participants with SCI had significantly larger max-MEPs in biceps maps than triceps maps (3.4±4.1 mV vs 0.5±0.3 mV, *p*<0.001), while able-bodied individuals had no differences in max-MEPs between biceps and triceps maps (0.8±0.6 mV vs 0.5±0.8 mV, *p*=NS). Notably, max-MEPs of the biceps map were abnormally higher in SCI than those observed in biceps maps of the control participants (*p*=0.02), even though max-MEPs of the triceps map did not differ between two groups of participants (*p*=NS).

**Figure 2.**
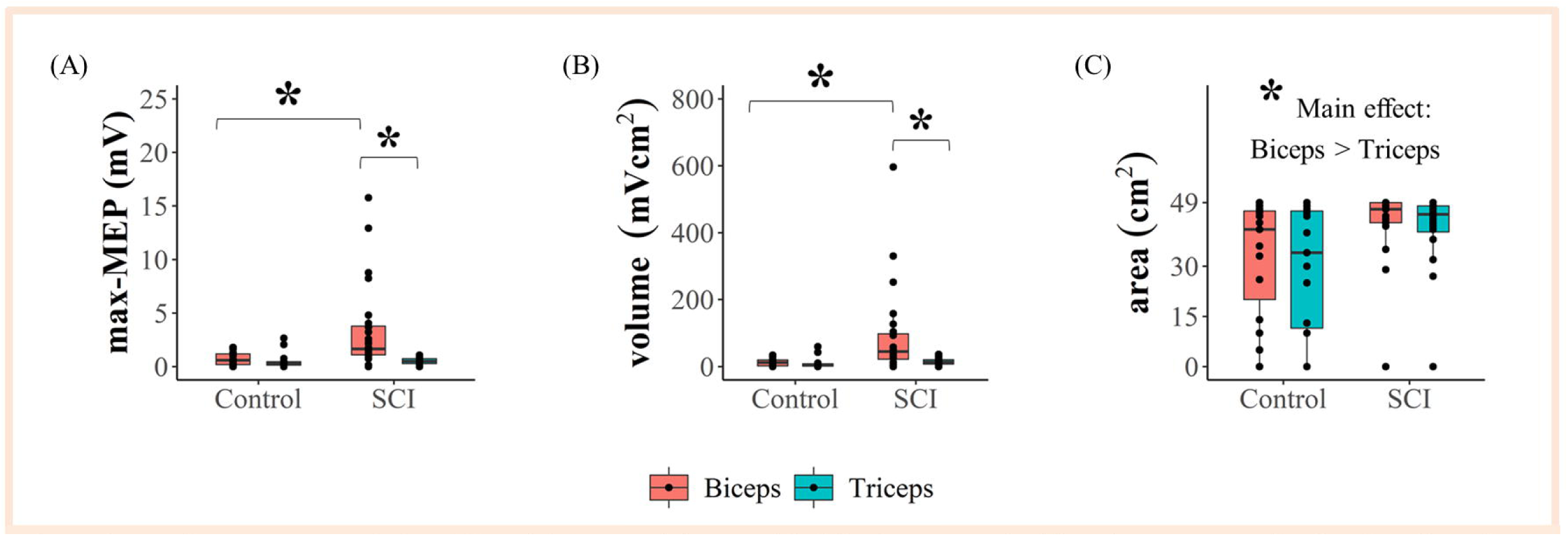
TMS map characteristics. **(A)** and **(B)** In participants with SCI, there were significantly greater maximal and total TMS map excitabilities devoted to the stronger biceps muscle than the weaker triceps muscle. Note that the maximal and total excitabilities of biceps maps in SCI were abnormally heightened even when compared with controls. **(C)** Biceps TMS maps showed larger areas than triceps TMS maps. *: p<0.05.

Total excitability of the TMS map defined using map volumes also demonstrated significant main effects of GROUP (F_(1,37)_=5.3, *p*=0.027) and MUSCLE (F_(1,37)_=5.4, *p*=0.025) and an interaction effect (F_(1,37)_=5.0, *p*=0.032, **Fig. 2B**). Posthoc analysis revealed that participants with SCI had significantly larger volumes in biceps maps than triceps maps (92.0±134.2 mV*cm^2^ vs 14.5±10.0 mV*cm^2^, *p*<0.001), while able-bodied individuals had no differences in volume values between biceps and triceps maps (12.4±11.0 mV*cm^2^ vs 10.5±17.3 mV*cm^2^, *p*=NS). Similar to that for max-MEPs, volumes of the biceps map were abnormally higher in SCI than those observed in biceps maps of the control participants (*p*=0.004). However, volumes of the triceps map did not differ between two groups of participants (*p*=NS).

Map area had a significant main effect of MUSCLE (F_(1,37)_= 8.0, *p*=0.008) but not of GROUP (F_(1,37)_= 3.2, *p*=NS) or interaction (F_(1,37)_=0.2, *p*=NS, **Fig. 2C**). TMS map area was larger for biceps than triceps with a mean (SD) difference of 3.5 (7.0) cm^2^ (*p*=0.004).

COG (x) had a significant main effect of MUSCLE (F_(1,31)_=11.0, *p=*0.002) and GROUP X MUSCLE interaction (F_(1,31)_= 4.9, *p=*0.03) but no main effect of GROUP (F_(1,31)_= 0.28, *p*=NS). Despite a significant interaction effect, Bonferroni-corrected posthoc analyses failed to reveal significant differences in COG (x) for either map between participants with SCI and able-bodied controls (*p*=NS). Biceps COG was slightly more lateral than triceps COG with a mean (SD) difference of 1.0 (2.0) cm (*p*=0.029). As to COG(y), there was a significant main effect of MUSCLE (F_(1,31)_= 21.9, *p*<0.01) but not of GROUP (F_(1,31)_<0.01, *p=*NS) or the interaction (F_(1,31)_= 3.65, *p=*NS). Biceps COG was slightly more anterior than triceps COG with a mean (SD) difference of 2.0 (2.4) cm (*p*<0.001).

### Correlations

Correlation analyses confirmed positive relationships between biceps and triceps MVCs and upper arm CUE-T scores. As one would expect, participants with SCI who had stronger MVC activation for the two muscles also exhibited better upper arm motor performance (*ρ*=0.82, *p*<0.001 and *ρ*=0.80, *p*=0.001, respectively).

Biceps and triceps map excitability measures were found positively associated with upper arm CUE-T (**Fig. 3**). Participants with greater max-MEPs of biceps maps (*ρ*=0.52, *p*=0.01; **Table 2**, **Fig. 3**) and max-MEPs (*ρ*=0.58, *p*=0.004) and volumes (*ρ*=0.44, *p*=0.03) of triceps maps (**Table 2**, **Fig. 3**) had greater upper arm motor function.

**Figure 3.**
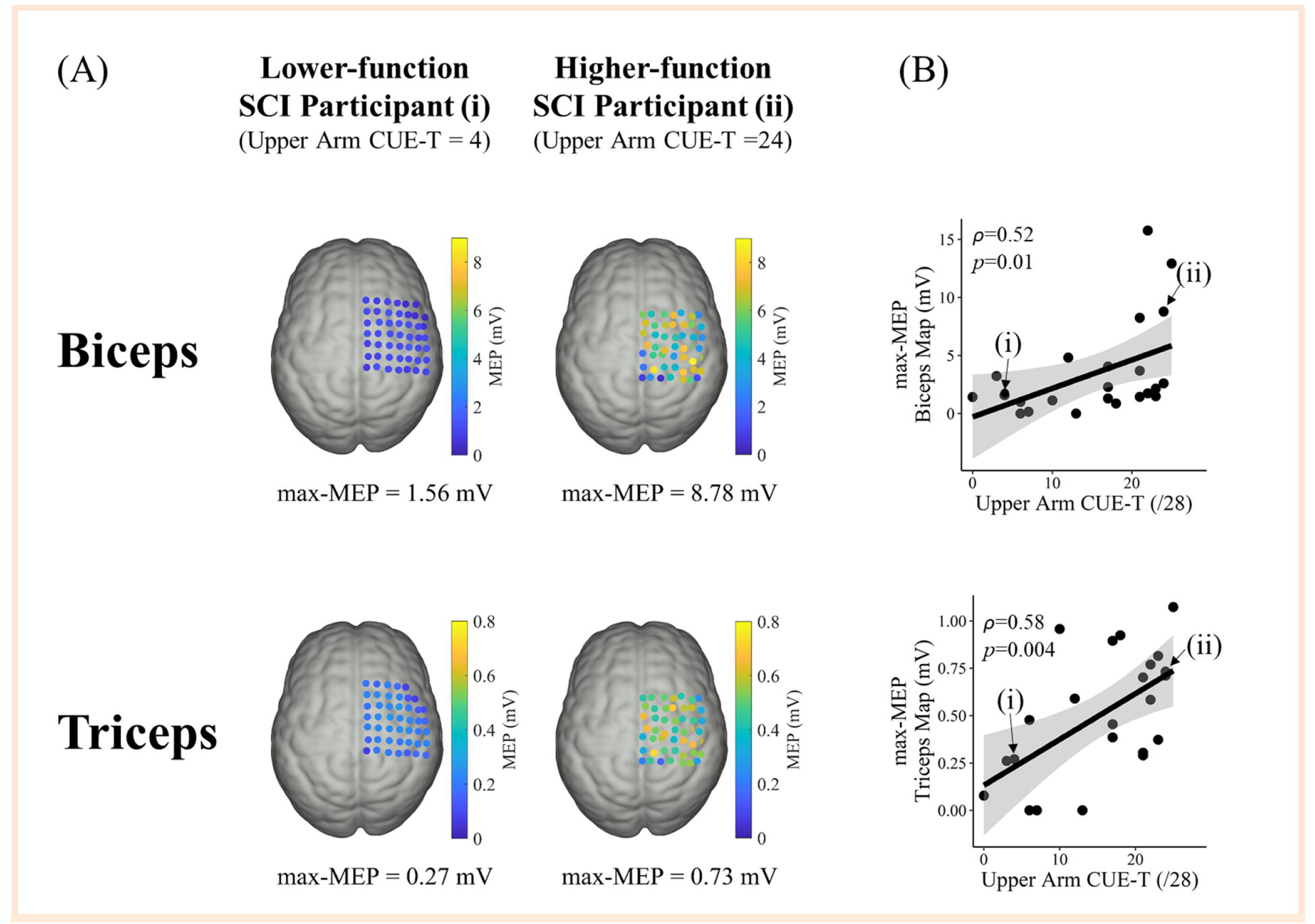
Relationships of TMS map max-MEPs with upper arm CUE-T scores. **(A)** Representative TMS maps of triceps and biceps muscles from a lower-function (left, indexed as i) and a higher-function (right, indexed as ii) participant with SCI. The color bar indicates MEP amplitudes of active sites in each TMS map. **(B)** Participants with higher motor function demonstrated greater maximal excitability in both of their triceps and biceps maps; and vice versa. Grey shades represent I standard error of the estimated correlation slope.

**Table 2.** Correlations of TMS map characteristics to upper arm CUE-T scores and MVCs.

|  | Upper Arm<br>CUE-T Score |  | MVC of<br>Homologous Muscle |  |
| --- | --- | --- | --- | --- |
| | $\rho$ | $p$ | $\rho$ | $p$ |
| **Biceps Map** |  |  |  |  |
| area | 0.13 | 0.56 | 0.21 | 0.34 |
| max-MEP | **0.52** | **0.01\*** | **0.73** | **<0.001\*\*\*** |
| volume | 0.31 | 0.15 | **0.57** | **0.004\*\*** |
| COG (x) | -0.07 | 0.78 | -0.01 | 0.97 |
| COG (y) | 0.10 | 0.67 | -0.03 | 0.90 |
| **Triceps Map** |  |  |  |  |
| area | 0.26 | 0.24 | 0.14 | 0.53 |
| max-MEP | **0.58** | **0.004\*\*** | 0.28 | 0.19 |
| volume | **0.44** | **0.03\*** | 0.18 | 0.40 |
| COG (x) | -0.01 | 0.97 | -0.25 | 0.27 |
| COG (y) | 0.19 | 0.42 | -0.10 | 0.67 |

TMS=transcranial magnetic stimulation; CUE-T=Capabilities of Upper Extremity Test; MVC=maximal voluntary contraction; MEP=motor evoked potential; COG=center of gravity. Bold number with \*: $p < 0.05$ , \*\*: $p < 0.01$ , \*\*\*: $p < 0.001$ .

While triceps MVCs and triceps map excitability were not related, biceps MVCs were positively correlated to biceps excitability (**Fig. 4**). Participants with higher biceps maximal activation showed larger biceps map max-MEPs (*ρ*=0.73, *p*<0.001; **Table 2**, **Fig. 4**) and volumes (*ρ*=0.57, *p*=0.004; **Table 2)**.

**Figure 4.**
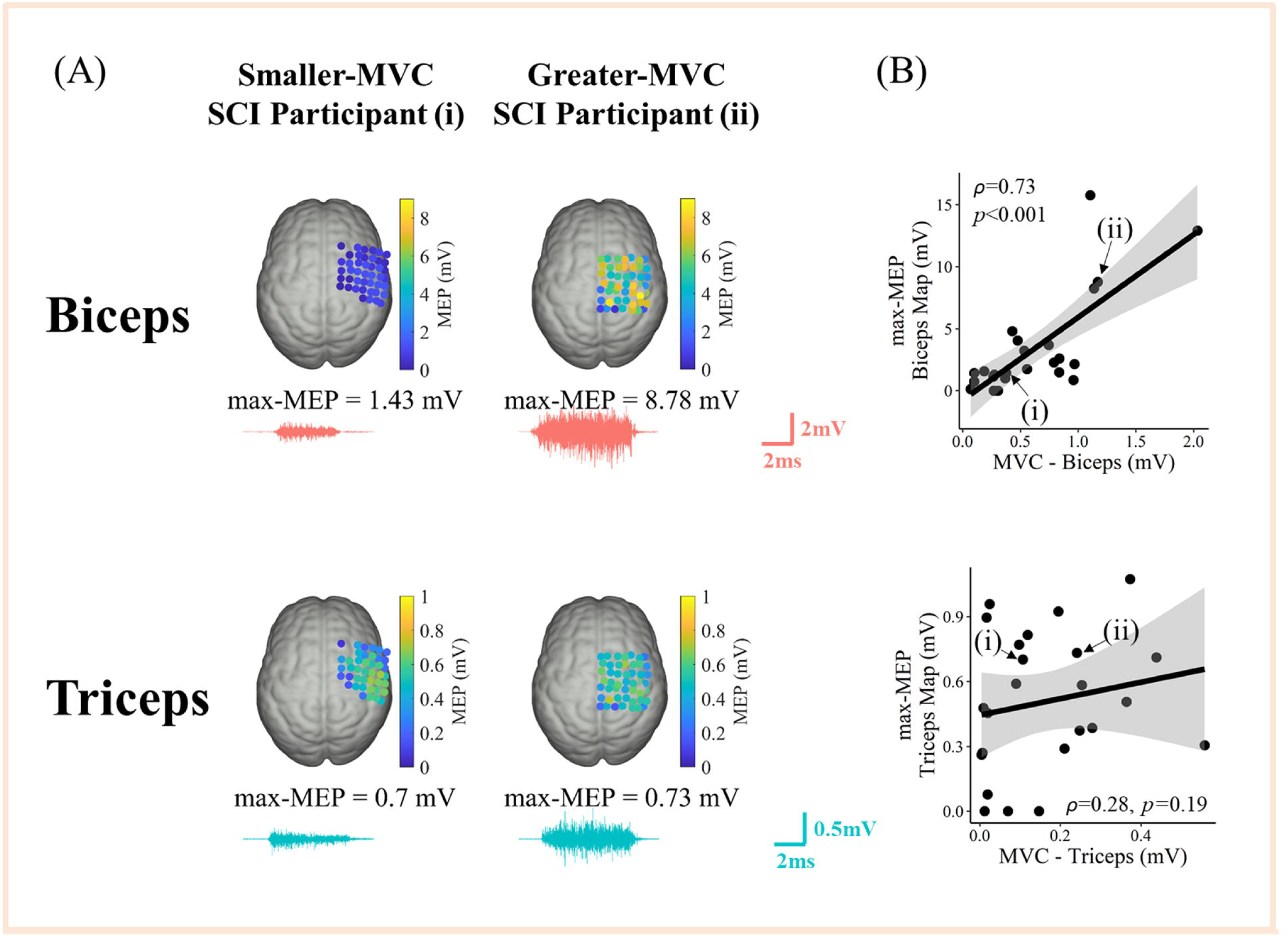
Relationships of TMS map max-MEPs with respective muscle’s MVCs. **(A)** Representative biceps and triceps TMS maps from two participants with SCI who demonstrated smaller MVCs (left, indexed as i) and larger MVCs (right, indexed as ii) of two muscles. The color bar indicates MEP amplitudes of active sites in each TMS map. **(B)** There was a positive relationship between TMS map max-MEPs and MVCs in the biceps muscle (top) but not the triceps muscle (bottom). Grey shades represent one standard error of the estimated correlation slope.

In addition, participants with SCI who had more distal injuries (i.e., lower NLIs, which would indicate more cervical spinal segments were spared) had higher excitability of biceps maps (max-MEP: *ρ*=0.47, *p*=0.02) and triceps maps (max-MEP: *r*=0.52, *p*=0.01; volume: *r*=0.49, *p*=0.02). Further, participants with higher map excitability of the weakened triceps muscle had more sensory sparing (i.e., more distal sensory levels, max-MEP: *ρ*=-0.51, *p*=0.014 and volume: *ρ*=-0.60, *p*=0.003) and less triceps muscle spasticity (max-MEP: *ρ*=-0.59, *p*=0.003 and volume: *ρ*=-0.57, *p*=0.005). There were no significant relationships between map excitability measures and SCI chronicity, AIS grades, or motor levels.

## Discussion

The purpose of this study was to evaluate characteristics of TMS motor maps of biceps and triceps muscles in persons with tetraplegia following chronic traumatic cervical SCI. We found that the biceps map is more excitable than the triceps map in participants with SCI and even more so than the biceps map of able-bodied control participants. Participants with SCI who had higher excitability of biceps and triceps maps also had better upper arm functional ability scores and more distal neurological level of injury (i.e., have more spared cervical segments). Our cross-sectional results in a relatively homogenous sample of chronic traumatic cervical SCI participants substantiated preclinical evidence of injury-induced deafferentation in SCI models, and for the first time revealed the relationship between reorganized motor cortical maps of muscles with different extents of sparing and upper arm motor status.

### Reorganization of Map Excitability but not Territory after SCI

Our results revealed that biceps maps were more excitable than triceps maps in participants with SCI. Notably, biceps maps in participants with SCI were also more excitable than biceps maps in able-bodied participants. Such hyper-excitable biceps representations in SCI have been reported as early as 6 days post-injury and known to persist into the chronic phase of SCI [6]. Muscle spasticity is unlikely to be contributing as our results revealed no correlation between biceps map excitability and biceps muscle spasticity. Other factors may provide more mechanistic insights. Heightened excitability in the acute stage has been attributed to unmasking of latent corticospinal neurons in response to SCI[6]. However, hyper-excitability in the chronic stage studied here is likely mediated through use- or activity-dependent plasticity [3]. Spared muscles tend to be used more in daily activities (at times to compensate for weaker muscles), which may invoke competition for neural resources and neurotrophic factors preconditioning neural regeneration and axonal sprouting[3]. The finding of biceps map excitability being associated with both MVCs of biceps and scores of upper arm functional tests supports use-dependent mechanisms at play in observed map plasticity.

Interestingly, triceps map excitability was not different between two groups of participants, even though triceps muscle was significantly weakened in SCI. The first potential explanation could be that motoneuron excitability at rest is abnormally heightened for affected muscles in humans with SCI [17] likely resulting from the reoccurrence of persistent inward currents and loss of supraspinal control [18]. However, it is unlikely that overactivity of motoneurons affected our results. Increased spinal excitability has been linked to muscle spasms [18], but our results revealed participants with greater triceps map excitability had less severe spasticity which is in agreement with findings of Sangari and Perez [19]. Another explanation is that sensory sparing likely helped maintain the tonic cutaneous inputs onto cortical and spinal motoneurons and thereby preserve corticomotor excitability even in weakened muscles. Previous studies reported that persons with sensory deficits showed reduced motor cortical map excitability [20, 21], which is in line with our results that participants with lower sensory levels (i.e., more preserved sensory segments) exhibited higher corticomotor excitability (measured by TMS map max-MEPs and volumes).

Over-excitability of biceps relative to triceps maps was not a function of larger cortical territory since there was no difference in map area across the two muscles in persons with SCI. Evidence of expanded motor map territory of segments that are spared and more proximal to injury site is not uncommon in deafferentation-based preclinical models [4]. Studies have revealed that representations of whiskers enlarge at the cost of forelimb representations in rodent models of SCI [4]. However, such across-representation shifts in territory are easier to observe when there are larger anatomic distances and distinct somatotopic gradients involved such as between whiskers and forelimbs [22]. But biceps and triceps muscles studied here naturally have abundant overlap in the motor cortex. COGs of biceps and triceps mostly overlapped and were only 1.0 cm and 2.0 cm apart at the scalp level in the medial-lateral and anterior-posterior directions, respectively. Post-hoc analyses revealed a median (25^th^ and 75^th^ percentiles) area overlap of 44.0 (40.3, 48.0) cm^2^ between biceps and triceps representations in participants with SCI. Anatomic proximity and overlap in motor networks therefore may explain why maps of biceps were not larger than maps of triceps in persons with SCI. Excitability or motor map activity measures may be more sensitive to identify reorganization in such cases of overlapping territories.

### Clinical Implications

Knowledge of clinical implications of neural plasticity is crucial for designing effective treatments. Our study, for the first time, demonstrated that motor map excitability of the relatively spared biceps muscle was associated with more favorable upper arm motor functional scores and maximal volitional activation of the muscle in SCI. Greater use of the stronger biceps muscle, and possibly shoulder girdle muscles, as a way to compensate for ineffective recruitment of weaker muscles in accomplishing daily activities and self-care tasks may explain heightened map excitability.

In addition, triceps map excitability was higher among participants who had better upper arm motor function. Such finding underscored the importance of residual excitability to weakened muscles in functional performance. Interestingly, unlike biceps, triceps map excitability was not related to maximal volitional activation of triceps studied using MVC.

Together, these results suggest that residual cortical map excitability may be implicated in tasks demanding submaximal but not maximal volitional effort, though longitudinal investigation of map changes and restoration (even if partial) of maximal volitional activation strength may offer a clear solution. Nevertheless, there lies an opportunity to identify adjunctive treatments that may help to elicit higher levels of activation from the more impaired muscle, whether through intensive use or neurostimulation such as transcranial direct current stimulation [23] or paired corticospinal-motoneuronal stimulation [24], and track gains in volitional muscle activation in association with map reorganization in chronic SCI survivors.

### Limitations

This study has a few limitations. First, we used 100% MSO to evaluate motor cortical maps of biceps and triceps muscles. The high intensity of TMS used here might have overestimated the size of muscle representations. However, our approach is consistent with that used in Brouwer and Hopkins-Rosseel [7] and Levy Jr et al [8] who previously evaluated and contrasted cortical presentations of biceps and triceps muscles in small samples of participants with cervical SCI (n=10 in [7] and n=2 in [8]). Second, we studied the motor maps at resting state. Cautions are needed when generalizing our results to studies of muscle representations evaluated at the active state. Third, correlations of motor map characteristics to upper limb function were studied at the baseline of a clinical trial. Longitudinal results are needed to understand changes of motor map amplitudes in response to treatment gains.

## Conclusion

In summary, our study revealed that motor cortical maps of spared biceps muscle became hyper-excitable than maps of the weaker triceps muscle in a relatively homogenous sample of persons with chronic traumatic cervical SCI. Use-dependent mechanisms resulting from typically over-recruitment of biceps muscle may be implicated though influence of altered spinal networks remains to be investigated. Higher map excitability in general was associated with better upper limb motor scores. While map excitability for biceps was also associated with maximal muscle activation in persons with SCI, map excitability for triceps was not suggested. Excitability of maps of weaker muscles may have a relationship with tasks requiring less-than-maximal volitional effort. There is an opportunity for motor map characteristics to serve as biomarkers of response to treatments aimed at augmenting upper limb rehabilitation in cervical SCI.

## Data Availability Statements

Data are presented within the article. Further data will be available for scientific communication upon request.

## Acknowledgments

The U.S. Army Medical Research Acquisition Activity, 820 Chandler Street, Fort Detrick MD 21702-5014 is the awarding and administering acquisition office. We would like to acknowledge the roles of all team members across study sites, including coordinators, therapists, research assistants, and engineers. We thank the research participants for their valuable time and efforts.

## Author contributions

**Conception or design of the work.** EP, JL, KB, KO, GF, TA, SK, SP, KK, AB, XW

**Acquisition, analysis or interpretation of data for the work.** JL, KO, EP, TA, DA, GN, FB, MKH, NB, MB

**Drafting the work or revising it critically for important intellectual content.** JL, EP, KO, GN, FB, SP, GF, SK, KK, AB, MKH, XW, KB, NB, MB

